# *In situ* generation of Aβ_42_ oligomers via secondary nucleation triggers neurite degeneration and synaptic dysfunction in human iPSC-derived glutamatergic neurons

**DOI:** 10.1101/2024.08.30.610591

**Authors:** Alicia González Díaz, Eleonora Sarracco, Andrea Possenti, Isaac Kitchen-Smith, Sean Chia, Joseph Menzies, Gabriel Stephenson, Rodrigo Cataldi, Kim Yahya, Yuqi Bian, Gustavo Antonio Urrutia, Sara Linse, Benedetta Mannini, Michele Vendruscolo

## Abstract

The aggregation of Aβ42 into misfolded oligomers is a central event in the pathogenesis of Alzheimer’s disease. In this study, we aimed to develop a robust experimental system that recapitulates Aβ42 oligomerization in living cells to gain insight into their neurotoxicity and to provide a platform to characterize the effects of inhibitors of this process. Our strategy is based on the *in situ* generation of Aβ42 oligomers via secondary nucleation by repeatedly treating the cells with Aβ42 monomers in the presence of pre-formed Aβ42 fibrils. This approach enables an accurate control over the levels of on-pathway soluble Aβ42 oligomers and cell-associated aggregates, as well as the study of their neurotoxic effects. By implementing this approach in human glutamatergic neurons derived from induced pluripotent stem cells (iPSCs), we were able to replicate key aspects of Alzheimer’s disease, including neurite degeneration and synaptic dysfunction. Using BRICHOS, a molecular chaperone that specifically inhibits secondary nucleation, we confirmed that aggregation in this system occurs through secondary nucleation, and that quantitative parameters for comparing potential Aβ42 aggregation inhibitors can be obtained. Overall, our results demonstrate that by *in situ* generation of on-pathway Aβ42 oligomers, one can obtain translational cellular models of AD to bridge the gap between basic research and clinical applications.

## Introduction

The misfolding and aggregation of Aβ into amyloid plaques is a histopathological hallmark of Alzheimer’s disease (AD)^1–6^. The nature of the involvement of Aβ in the pathological processes of AD has therefore been the object of intense research^4–6^. Aβ results from the amyloidogenic processing of the amyloid precursor protein (APP) by β-and ɣ-secretases^4–6^. Mutations in *APP*, as well as *PSEN1* and *PSEN2*, which are genes encoding components of the γ-secretase complex, are characteristic of familial AD (fAD)^4–6^. By contrast, age-related changes in the neuroinflammatory and degradation pathways and perturbations of the protein homeostasis system are typical of sporadic AD (sAD)^4–6^. These are all factors that can exacerbate the production, hinder the clearance, and facilitate the aggregation of Aβ peptides in the brain^4–6^.

As the extent of the deposition of Aβ aggregates into amyloid plaques does not correlate with the disease severity and the dysregulation of Aβ is a marker characteristic of early disease stages, it has been suggested that the neurotoxic effects of Aβ aggregation are caused by small soluble aggregates preceding fibril formation and referred to as oligomers^4–10^ (**Figure 1A**). Aβ oligomers exist in the brain as structurally heterogeneous and transient species, and their mechanisms of toxicity depend on their biophysical properties, such as hydrophobicity, size and β-sheet secondary structure content^11,12^. Aβ oligomers have been reported to by cytotoxic through a variety of mechanisms, including by: (i) disrupting cell membranes, (ii) triggering reactive oxygen species (ROS) production, calcium influx, mitochondrial disruption and apoptosis, (iii) interacting with cell membrane receptors, and (iv) causing inflammatory responses, which can eventually lead to synaptic dysfunction and neurodegeneration^7–11^.

**Figure 1.**
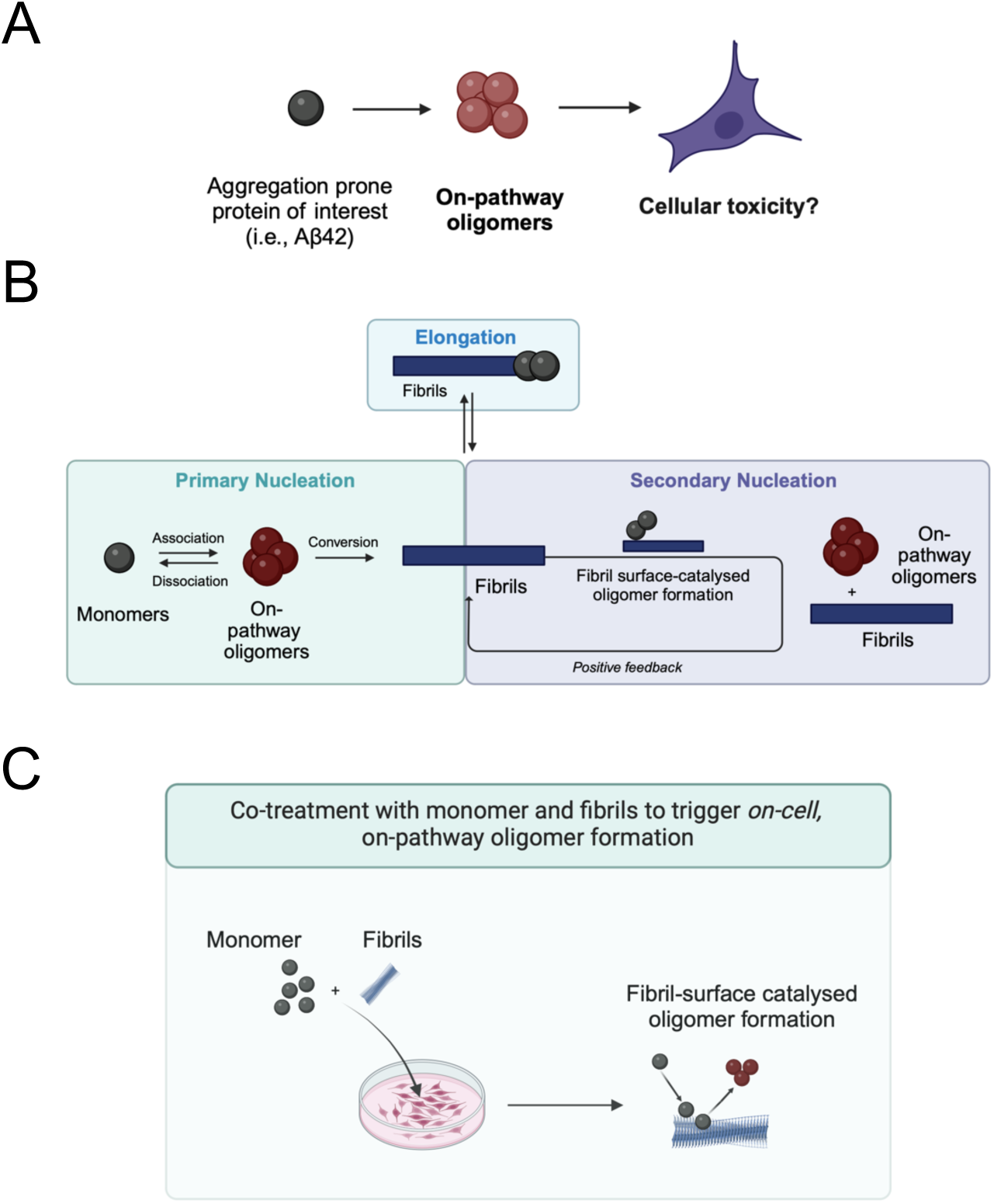
Summary of the approach reported in this work to recapitulate on-pathway Aβ_42_ oligomer generation and derived cellular toxicity. **(A)** Schematic representation of the hypothesis that Aβ_42_ oligomers generated on-pathway during the aggregation process are cytotoxic. **(B)** Overview of the kinetic mechanisms driving the conversion of Aβ_42_ into oligomers and fibrils. Via macroscopic measurements, an aggregation reaction of Aβ_42_ can be followed over time by tracking the total mass of fibrillar aggregates. Time points can be collected (t_lag_, t_half_ or t_plateau_) to examine the levels of discrete populations on-pathway oligomers. **(C)** Illustration showing the approach described in this work involving *in situ* generation of on-pathway Aβ oligomers to study their cytotoxic profile, without selecting any population from an *in vitro* kinetic reaction.

Given the relevance of the early phases of Aβ aggregation, Aβ oligomers have been investigated as a target in drug discovery^13,14^. The first disease-modifying treatments approved by the FDA for AD, aducanumab^15,16^, lecanemab^17,18^ and donanemab^19,20^, target Aβ aggregates and are likely to reduce the production of Aβ oligomers^16^. To continue the development of increasingly more effective drugs targeting these species, it is now important to address two challenges: the first is understanding of the mechanisms of formation of Aβ oligomers in a disease-relevant environment, and the second is develop systems that recapitulate the pathological features associated with AD (**Figure 1B,C**).

The mechanism of Aβ oligomer generation has been studied *in vitro* by using chemical kinetics^9,21,22^. The study of the *in vitro* aggregation process of Aβ has allowed to model the conversion of Aβ monomers into amyloid aggregates through a series of microscopic steps governed by specific rate laws^9,23^. These microscopic steps fall into two main categories: (i) those that increase the total number of aggregates, including primary and secondary nucleation, and (ii) those that influence the overall mass of the aggregates and their growth, such as fibril elongation and monomer dissociation (**Figure 1B**)^21,24^. Secondary nucleation is the process that most impact oligomer generation. In this process, the surface of existing amyloid fibrils catalyses the formation of new oligomers, establishing a positive feedback loop that generates increasing amounts of Aβ oligomers^21^. The *in vitro* chemical kinetics have been used to assess the potency of inhibitors of specific microscopic steps in Aβ aggregation^16,25–27^. However, this type of studies does not provide direct information on the impact that the aggregation process of Aβ and its modulation have on the functionality of the cells.

To this aim, we developed a robust cell assay to monitor the aggregation of Aβ_42_, the 42-residue form of Aβ, the oligomer production, and the phenotypic alteration associated with these processes. This approach combines the understanding from chemical kinetics that secondary nucleation is the main mechanism of formation of on-pathway oligomers^9,21,22^ and physiologically-relevant readouts to assess the levels of cellular dysfunction. To generate *in situ* Aβ_42_ oligomers through secondary nucleation, we promote Aβ_42_ aggregation by co-treating the cells with Aβ_42_ monomers and pre-formed Aβ_42_ fibrils (**Figure 1C**). By enforcing this aggregation mechanism directly onto the cells, we make it possible to evaluate the biological impact of all the on-pathway oligomeric species generated during the aggregation reaction. We show that the oligomers are generated by secondary nucleation by treating the cells with BRICHOS, a molecular chaperone shown to inhibit specifically this microscopic step^28^. We thus anticipate that this assay will enable the reliable testing of candidate inhibitors of Aβ_42_ aggregation, including inhibitors of primary or secondary nucleation, and the evaluation of the extent of the phenotypic recovery following the inhibition of the aggregation.

## Results

### Optimization of a protocol for the *in-situ* generation of Aβ_42_ oligomers in cholinergic-like SH-SY5Y cells

Our initial step was to develop and optimise a protocol to investigate the formation and biological activity of the Aβ_42_ oligomers generated during an aggregation reaction in a cellular system, without the bias of selecting or artificially stabilizing a particular population. For this purpose, we triggered *in situ* the secondary nucleation of Aβ_42_ soluble oligomers in cultured cells.

To carry out the optimisation on a robust cell system, we treated SH-SY5Y (human neuroblastoma) cells, differentiated into a cholinergic phenotype, either with Aβ_42_ monomers alone or with a combination of Aβ_42_ monomers and a low concentration of Aβ_42_ fibrils (**Figure 2**). The comparison of these two approaches was motivated by *in vitro* aggregation studies that indicated that Aβ_42_ monomers, in the absence of fibrils, aggregate via a primary nucleation mechanism to form oligomers and amyloid fibrillar species^23,24^. These fibrils can then accelerate the formation of Aβ_42_ oligomers via a secondary nucleation process^21,24^. Therefore, with our comparison we aimed to study the Aβ aggregation process in the presence of the cells, both driven by Aβ_42_ monomers alone and by Aβ_42_ monomers and fibrils, which could catalyse and accelerate *in situ* Aβ_42_ oligomer generation.

**Figure 2.**
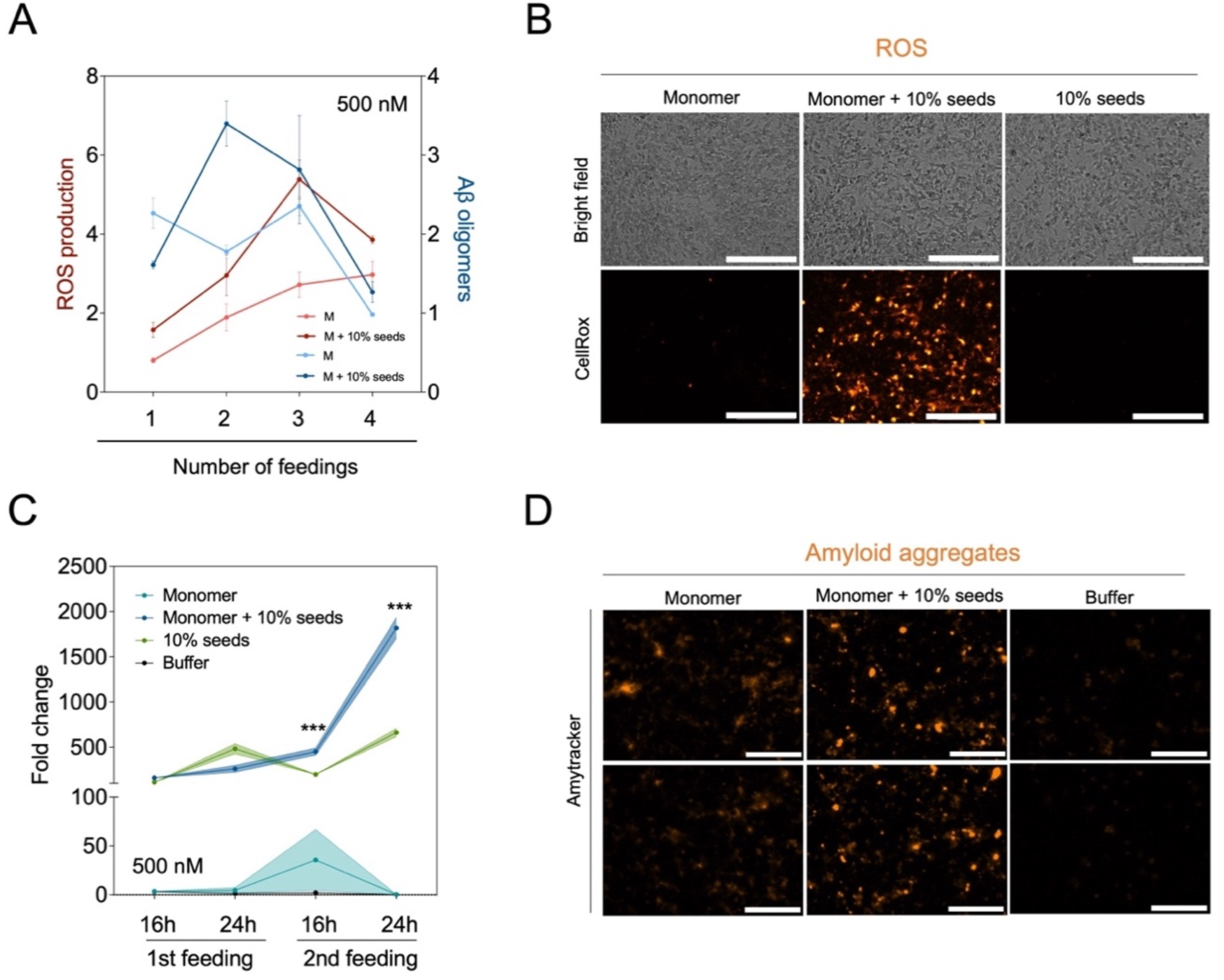
The number of co-treatments and the concentrations of Aβ_42_ monomers and pre-formed Aβ_42_ fibrils impacts the Aβ_42_ oligomerisation window and concurrent ROS in cholinergic-like human SH-SY5Y cells. **(A)** Cholinergic-like SH-SY5Y cells were treated every 24 h for 4 consecutive days with 500 nM monomeric Aβ_42_ and 50 nM of pre-formed fibrils (fibrils). 24 h after each treatment, soluble Aβ_42_ aggregates (oligomers) were quantified from the supernatant of treated cells by a 6E10-6E10 homotypic ELISA and, concurrently, cells were stained with CellRox to assess ROS production via live-cell imaging. Data are reported as fold change of the total absorbance (Aβ_42_ oligomer ELISA) or normalised fluorescence (ROS) over the cells treated only once with buffer (n = 5). **(B)** Representative images of the CellRox-derived fluorescence on cholinergic-like SH-SY5Y cells treated twice with 500 nM of Aβ_42_ monomer, 500 nM of Aβ_42_ monomer (monomer) and 50 nM of preformed Aβ_42_ fibrils (monomer + 10 % fibrils), or 50 nM of pre-formed Aβ_42_ fibrils (fibrils) Scale bar = 200 µm. **(C)** 16 h and 24 h after 1 or 2 treatments (feedings) with either 500 nM of Aβ_42_ monomer or 500 nM of monomer and 50 nM of preformed fibrils, cells were fixed and stained with Amytracker to quantify the levels of amyloid aggregates^44^. Data are reported as fold change of the total Amytracker positive area over the cells treated only with monomer. Statistical differences were calculated by applying a two-way ANOVA analysis, using the Tukey’s test to correct for multiple comparisons (n=5, ***p-value < 0.001). **(D)** Representative images of Amytracker-positive signal on cholinergic-like SH-SY5Y cells treated twice with 500 nM of Aβ_42_ monomer, 500 nM of Aβ_42_ monomer (monomer) and 50 nM of preformed Aβ_42_ fibrils (monomer + 10 % fibrils) or buffer (no protein). Scale bar = 100 µm.

We treated the cells for four times every 24 h with 500 nM of Aβ_42_ monomers, in the absence or presence of pre-formed Aβ_42_ fibrils at 10% of the monomer concentration (50 nM). To follow the aggregation reactions, after each treatment we measured the concentration of soluble Aβ_42_ oligomers in the supernatant using ELISA (see Methods) and, simultaneously, assessed the production of reactive oxidative species (ROS) (**Figures 2A** and **S1**). We found that two repeated treatments were needed to obtain the optimal window in Aβ_42_ oligomer levels between seeded and non-seeded treatments. Additional treatments resulted in lower levels of detectable soluble Aβ_42_ oligomers in both conditions, likely due to Aβ_42_ aggregates maturing and getting trapped within cells and cell membranes (**Figure 2A**). ROS production was higher under seeded, compared to non-seeded, conditions at all the time points tested, indicating that aggregation products of Aβ_42_ induced by the fibrils trigger higher cellular oxidative stress (**Figure 2A,B**). To detect not only the soluble species but also those bound or internalized into the cells, we treated cholinergic-like SH-SY5Y twice with Aβ_42_ monomers in the absence or presence of fibrils, and quantified the aggregates by staining with Amytracker, an amyloid binding dye, at two time points, 16 h or 24 h after each treatment (**Figure 2C,D**). We refer to the species detected by Amytracker as cell-associated aggregates, as the staining protocol applied does not allow the detection of soluble species, which are washed away when the excess of dye is removed. After two exogenous treatments, we observed a significant increase in the levels of aggregates only in the cells co-treated with Aβ_42_ monomer and fibrils. As a control, cells treated with fibrils alone did not show such an increase, indicating that the aggregates observed upon co-treating with monomer and fibrils were the result of the fibril-catalysed aggregation reaction. By contrast, a significant increase in amyloid aggregates could not be observed in the cells treated with Aβ_42_ monomers over the time points assayed, indicating that the species forming under these conditions are of lower abundance and less competent to bind Amytracker (**Figure 2C,D**).

Taken together, these observations suggest that, in a cellular environment, the aggregation of Aβ_42_ monomers in the presence of pre-formed Aβ_42_ fibrils results in a greater amount of both soluble and cell-bound aggregates, compared to the aggregation of Aβ_42_ monomers without fibrils. Such differences in the aggregation process led to a diverse capacity of each treatment type to cause cellular dysfunction, with fibril-mediated Aβ_42_ aggregation triggering higher levels of ROS.

The experimental setup described above allowed us to achieve control and reproducibility on the Aβ_42_ aggregation onto the cells and opened the possibility of reliably testing inhibitors of Aβ_42_ on-pathway aggregation. The protocol is the result of an optimization that took into consideration the percentage of fibrils used to induce the aggregation of the monomer, the concentration of monomer, the number of treatments, the cell type, the readouts to detect aggregates and the readouts to assess the phenotypical dysfunction. The key experiments to set up this protocol are reported in **Figure S1**. Firstly, we addressed the optimal concentration of fibrils by performing a titration of fibril concentrations raging from 1% up to 10% relative to the monomer concentration fixed at 500 nM on non-differentiated SH-SY5Y cells. The addition to Aβ_42_ monomers of increasing concentrations of Aβ_42_ fibrils resulted in a dose-dependent increase in the amount of Aβ_42_ aggregates and ROS levels, compared to cells treated solely with equivalent Aβ_42_ monomer or fibrils concentrations (**Figure S1A,B**). We chose the concentration of 10% fibrils relative to the monomer as this was able to give the best window among the different tested conditions.

Secondly, to improve the extent of signal and reproducibility of the phenotypical readout, we explored the approach of enhancing the vulnerability of the cell system by differentiating the SH-SY5Y cells to cholinergic-like neurons (**Figure S1C-E**). The differentiated cells displayed more pronounced sensitivity to the Aβ_42_ exogenous treatment as compared to their undifferentiated counterparts in three different phenotypical readouts of cell dysfunction: ROS production (**Figure S1C**), chromatin condensation (**Figure S1D**) and calcium influx (**Figure S1E**). Overall, the co-treatment with Aβ_42_ monomers and fibrils resulted in increased cellular dysfunction as compared to cells treated solely with Aβ_42_ monomer or buffer.

We then investigated the optimal monomer concentration and number of treatments. We performed these experiments on cholinergic-like SH-SY5Y cells by tuning the Aβ_42_ monomer concentration from 500 nM to 4 µM, while keeping the concentration of pre-formed Aβ_42_ fibrils at 10% relative to Aβ_42_ monomers. The treatment regime was kept for four days and repeated every 24 h. After each treatment, soluble Aβ_42_ oligomers and ROS levels were measured (**Figures 2A** and **S1F-H**). While we could observe a higher level of ROS in the seeded condition compared to the non-seeded at all monomer concentration and number of treatments, we found that the best window to observe oligomerization was at the lowest monomer concentration, 500 nM, and after two treatments. Higher Aβ_42_ monomer levels (ζ 1 µM) and repeated treatments over two times led to comparable levels of soluble Aβ_42_ aggregates both in seeded and unseeded conditions (**Figures 2A** and **S1F-H**).

This experimental setup makes it possible to robustly recapitulate in a cellular system the cytotoxicity of on-pathway Aβ_42_ aggregation. For comparison, we tested another approach reported in the literature for the same purpose^22^. This approach consists in the exogenous treatment of cells with Aβ_42_ aggregates collected from an *in vitro* aggregation reaction conducted in the presence of seeds (**Figure S2A**). We assessed three time points of the aggregation kinetics: (i) before adding the seeds to the Aβ_42_ monomers (t_0_), (ii) in the lag-phase (t_lag_) 10 min after adding the seeds, and (iii) at the half-time (t_half_), 40 min after adding the seeds. Each stage is populated by different Aβ_42_ species: t_0_ is enriched with monomeric Aβ_42_, while t_lag_ and t_half_ samples progressively contain more on-pathway Aβ_42_ oligomers.^22^ SH-SY5Y cells were treated with 1 µM (monomer equivalents) of t_0,_ t_lag_ and t_half_ species to investigate both early (30 min to 1 h post-treatment) and late (24 h post-treatment) effects on ROS production and calcium influx (**Figure S2**). In our hands, this protocol resulted in a substantial variability among replicates in the toxicity profiles tested. The samples at t_0_ and t_lag_ showed no significant effect in increasing ROS (**Figure S2A,B**) or intracellular calcium levels (**Figure S2C,D**), as compared to buffer treated cells for both treatment regimes. By contrast, the samples at t_half_ induced, at least for some replicates, higher levels of ROS production both at early and late time points (**Figure S2A,B**). However, no significant differences in the calcium influx were observed between treatment groups (**Figure S2C,D**). Overall, these findings suggest that using samples directly extracted from an Aβ_42_ *in vitro* aggregation reaction to study aggregate-derived cytotoxicity lacks robustness, even for the samples enriched in Aβ_42_ oligomers (t_half_). This result is likely due to the metastable nature of the on-pathway Aβ_42_ aggregates and the consequent difficulty in isolating them. This method, in addition, has the disadvantage of allowing the test on cells only of a selection of populations of aggregates collected from the aggregation reaction *in vitro*.

### The *in situ* generation of Aβ_42_ oligomers recapitulates neurite degeneration and synaptic impairment in human iPSC-derived glutamatergic neurons

To improve the translational power of AD models based on the *in situ* Aβ_42_ oligomer generation protocol described above, we applied it to human iPSC-derived iCell glutamatergic neurons (see Methods). The neurons were cultured for 28 days and then treated twice with either 250 nM or 500 nM of Aβ_42_ monomers in absence or presence of 10% pre-formed Aβ_42_ fibrils (**Figure 3A,B**). After this repeated treatment, we analyzed first the levels of Aβ_42_ aggregates bound to the cells using Amytracker (**Figure 3C,D**). Both in the unseeded and seeded reactions, the increase of concentration of Aβ_42_ monomers from 250 nM to 500 nM promoted a progressive increase in the formation of aggregates, as shown by the quantification of the total area covered by Aβ_42_ aggregates positive to Amytracker. At both monomer concentrations, the seeded reaction led to a significant increase in aggregate levels compared to the unseeded reaction (**Figure 3D**). Since the contribution to the total Amytracker-positive area in neurons treated solely with fibrils is negligible, it is likely that these aggregates were primarily formed through *in situ* fibril-mediated aggregation of Aβ_42_ monomers. Aggregates were also detected under unseeded conditions. Although the total Amytracker-positive area was significantly lower under unseeded conditions than under seeded conditions, it was still significantly higher than in untreated neurons and those treated only with seeds at the highest tested monomer concentration (500 nM). Furthermore, seeding Aβ_42_ aggregation in cells influenced not only the maturation and ability to bind Amytracker, but also the morphology of the resulting aggregates (**Figure S3**). Specifically, cells treated with both Aβ_42_ monomers and fibrils exhibited brighter aggregate species with a denser core and protruding spikes. In contrast, repeated dosage with only Aβ_42_ monomers generated dim spiky aggregates spread across the cell surface (**Figure S3**).

**Figure 3.**
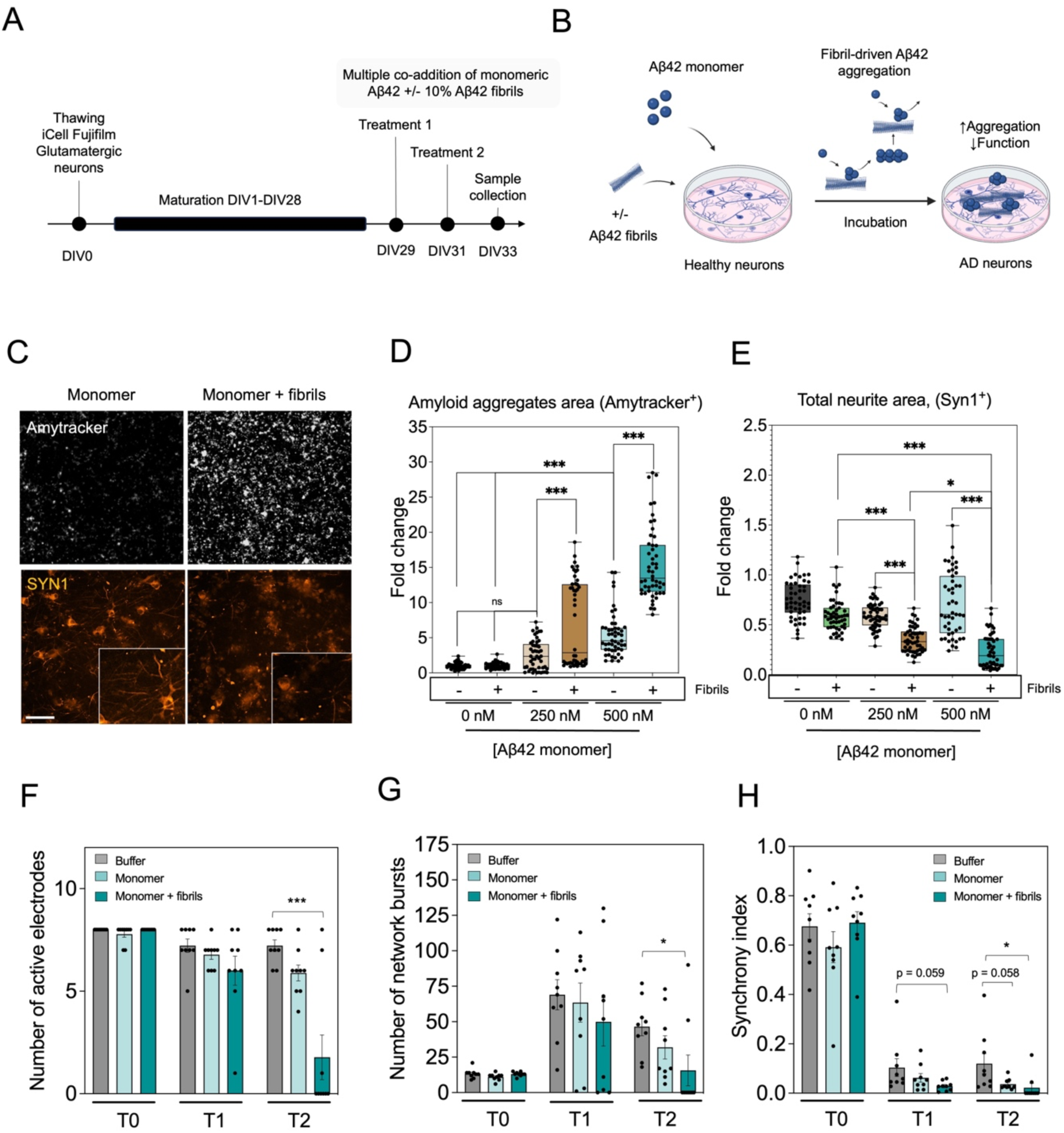
Multiple co-treatments of monomeric Aβ_42_ and pre-formed Aβ_42_ fibrils catalyse Aβ_42_ oligomer formation and triggers synaptic dysfunction in human iPSC-derived glutamatergic neurons. **(A)** Schematic representation of the experimental timeline. iCell glutamatergic human neurons were differentiated for 28 days *in vitro* (DIV). On DIV29 and DIV31, neurons were treated either with Aβ_42_ monomer, fibrils or monomer with fibrils at 10% of the monomer concentration. 48 h after the second treatment, samples were processed. **(B)** Figure representing how co-treating neurons with monomer and fibrils may promote secondary nucleation processes that drive the generation of new oligomeric species responsible for cellular dysfunction. **(C)** Representative pictures of the levels of Amytracker-positive aggregates and synapsin-1-positive glutamatergic neurons treated twice with 500 nM of monomeric Aβ_42_ or 500 nM Aβ_42_ and 50 nM of pre-formed fibrils. Scale bar = 100 µm. **(D,E)** Total area of Amytracker-positive aggregates (D) and total area of neurites positive for the synapsin-1 (Syn1) maker (E) of iCell glutamatergic neurons treated twice, every 48 h, with either seeded or non-seeded Aβ42 at concentrations of 250 nM or 500 nM. Each data point corresponds to the total area of amyloid aggregates or Syn1-positive signal per analysed picture. A total of 48-50 images were analysed per condition. Data are represented as fold change with respect to cells treated with buffer. Statistical differences were calculated by applying a one-way ANOVA analysis, using the Tukey’s test to correct for multiple comparisons (***p-value < 0.001). **(F-H)** Comparison on the electrophysiological activity profile of iCell glutamatergic neurons treated once (T1) or twice (T2) with buffer, 500 nM of Aβ_42_ monomer or 500 nM of monomer and 50 nM fibril, on a MEA plate. The number of active electrodes (F), number of network bursts (G) and synchrony index (H) are depicted (n = 9, mean±SEM). T0 represents the electrophysiological activity parameters from the corresponding MEA wells before they were treated with buffer, monomer or monomer + fibrils, respectively. Statistical differences were calculated by applying a one-way ANOVA analysis per treatment group, using the Dunnett’s test to correct for multiple comparisons (*p-value < 0.033, ***p-value < 0.001).

We then characterize the impact of these aggregation reactions onto the neurons by evaluating the integrity and functionality of the neuronal network by staining synapsin-1 (Syn1), a marker for synapses, and by measuring the electrophysiological activity. Glutamatergic neurons co-treated with Aβ_42_ monomer and fibrils showed clear morphological degeneration in their neurites compared to neurons treated with monomer only (**Figure 3C,E**). The quantification of the area of Syn1-positive neurites revealed a significant reduction in neurons subjected to seeded aggregation compared to those treated only with fibrils, at both monomer concentrations, indicating that the seeded aggregation has a significant impact in the synapse loss. There was no significant difference in this parameter between cells treated only with Aβ_42_ monomers and those treated only with fibrils (**Figure 3E**).

When we investigated the functionality of the neuronal network using the multi-electrode array (MEA) system (see Methods), we found that lower Syn1 levels correlated with impaired electrophysiological activity in the neurons treated with seeded monomer, with a drop in the number of active electrodes (i.e. detecting neuronal spikes), in the total levels of neuronal bursts and in the synchrony index (**Figure 3F-H**). Only using this assay we could detect a decrease in the synchrony index also in the neurons subjected to a repeated dosage only with monomer (**Figure 3F-H**). This observation was possible due to the higher sensitivity of the MEA technique as compared to the immunocytochemistry detection of Syn1, which was not able to reveal synaptic alterations for cells treated solely with the monomeric peptide.

Overall, these data show that the *in-situ* generation of Aβ_42_ oligomers recapitulates key hallmarks of the early phases of the AD pathology, including neurite degeneration and synaptic impairment in human iPSC-derived glutamatergic neurons.

### BRICHOS decreases Aβ_42_ aggregation in human iPSC-derived glutamatergic neurons upon co-treatment with Aβ_42_ monomer and fibrils

The findings reported above indicate that the seeded aggregation performed by co-treating the cells with Aβ_42_ monomers and pre-formed fibrils significantly increases the levels of both soluble oligomers and cell-associated aggregates. These results are in agreement with *in vitro* studies that show that the aggregation of Aβ_42_ catalysed by high-molecular-weight Aβ_42_ fibrillar species effectively generates the oligomers through a secondary nucleation process^21,24^. To confirm that the Aβ_42_ *in situ* oligomerization observed in the seeded aggregation on cells is governed by secondary nucleation, we tested BRICHOS in the experimental setup shown in **Figure 4**. BRICHOS, a human molecular chaperone, can specifically inhibit secondary nucleation by binding to the surfaces of fibrils, and redirecting the aggregation reaction to a pathway that minimizes the formation of oligomeric intermediates^28^. iCell Glutamatergic neurons were cultured for 28 days and then treated twice with 500 nM Aβ_42_ monomers in the presence or absence of 50 nM Aβ_42_ fibrils and 1 µM BRICHOS (**Figure 4**). We found that BRICHOS significantly decreased the levels of Aβ_42_ aggregates, including both soluble Aβ_42_ oligomers in the supernatant (**Figure 4C**) and aggregates bound to the cells (**Figure 4D,E**), as assessed by ELISA and staining with Amytracker, respectively.

**Figure 4.**
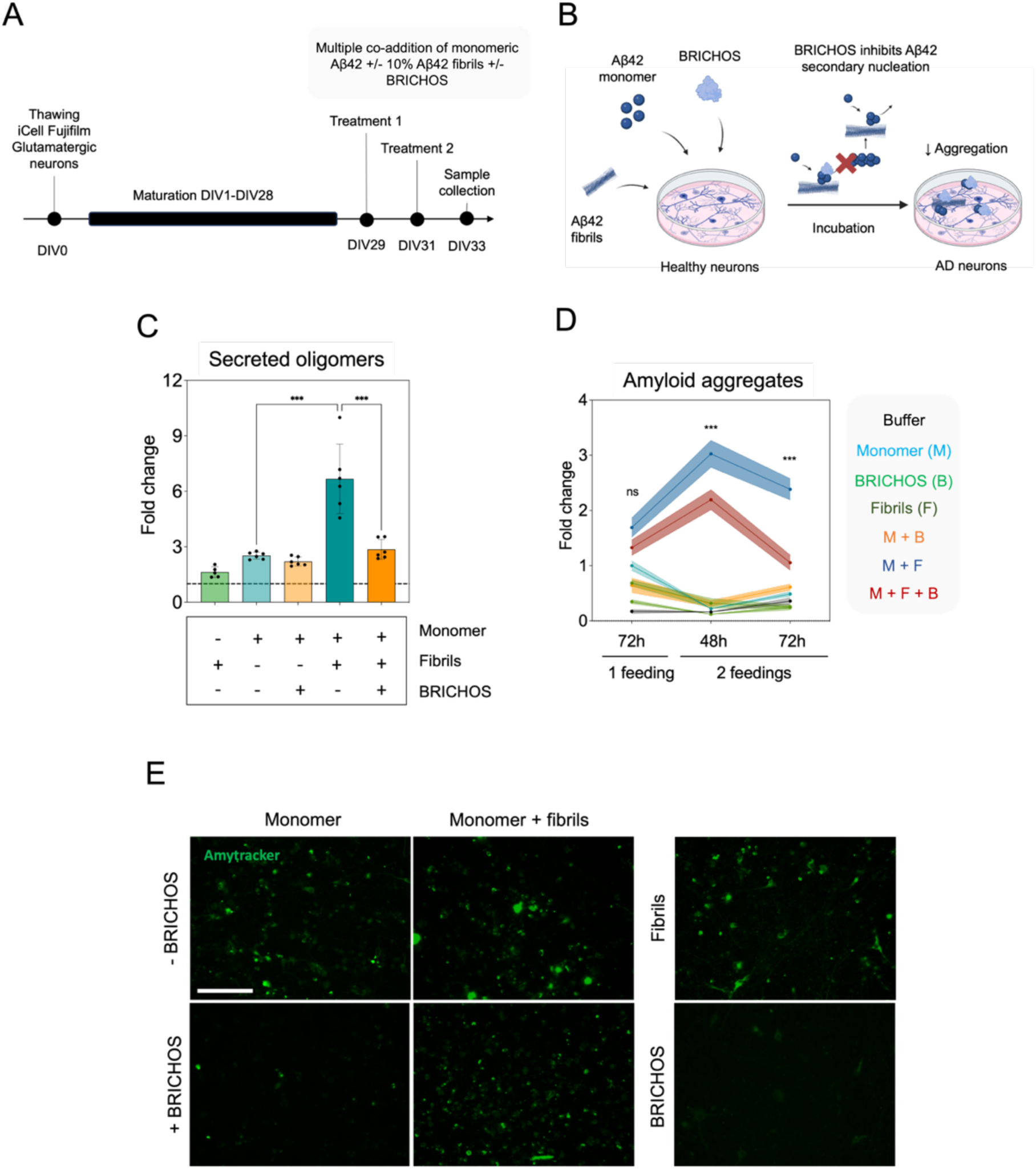
The secondary nucleation inhibitor BRICHOS decreases the levels of soluble Aβ_42_ oligomers and membrane-associated aggregates in glutamatergic neurons co-treated with Aβ_42_ monomers and pre-formed Aβ_42_ fibrils. **(A)** Schematic representation of the experimental timeline. iCell human glutamatergic neurons were kept in culture for 28 days. On DIV29 and DIV31, neurons were treated either with or without 1 µM of BRICHOS combined with: (i) 500 nM monomer, (ii) 50 nM fibrils or (iii) 500 nM monomer and 50 nM fibrils. 48 h after the second treatment, samples were processed. **(B)** Figure representing the addition of BRICHOS as an approach to inhibit or reduce aggregation events driven by secondary nucleation processes. **(C)** Quantification of the levels of soluble Aβ_42_ aggregates (oligomers) on the supernatant of glutamatergic neurons treated with monomer and/or fibrils in the presence or absence of BRICHOS, by a 6E10-6E10 homotypic ELISA. Data are reported as fold change of the absorbance over the cells treated with buffer (n = 6). **(D)** Progression of the total area of Amytracker-positive aggregates of neurons treated once or twice with seeded or non-seeded Aβ_42_ in the presence or absence of BRICHOS 72 h after the first feeding. The data are represented as the mean and standard error of the fold change of the levels of amyloid aggregates with respect to cells treated only with buffer. A total of 25-50 images were analysed per condition and time point. buffer. Statistical differences were calculated by applying a two-way ANOVA, using the Tukey’s test to correct for multiple comparisons (ns p-value > 0.12, ***p-value < 0.001). **(E)** Representative images of Amytracker-positive aggregates of treated neurons 72 h after the second treatment (scale bar = 100 µm).

These data confirm that our protocol can accurately replicate Aβ_42_ aggregation and the generation of on-pathway oligomers, which are key events in the onset and progression of AD. Additionally, the observed effects of BRICHOS suggest that this system may be used to screen other candidate compounds targeting Aβ42 oligomers and to study target engagement, which highlights its translational potential.

### Determination of quantitative parameters for inhibitors of Aβ_42_ oligomer formation

As next step, we modified the experimental setup to enable the quantitative investigation and comparison of the effects of Aβ_42_ aggregation inhibitors. To this aim, we measured the Aβ_42_ aggregation time course directly on the neurons using fluorescently labelled Aβ_42_ monomer (**Figure 5A**). A strategy based on *in vitro* chemical kinetics has been used for rational drug discovery to analyse quantitatively the effects of small molecules on the rates of specific microscopic steps in Aβ_42_ aggregation^29^. In this modified experimental design, iCell glutamatergic neurons were treated with 250 nM and 500 nM of fluorescently-labelled Aβ_42_ monomers in the presence or absence of 10% pre-formed unlabelled Aβ_42_ fibrils and the aggregation reactions were monitored over time (**Figure 5A)**. The treatment with Aβ_42_ monomers alone at both concentrations did not show any increase in the number of aggregates segmented as bright objects over the recorded time. By contrast, neurons co-treated with 500 nM Aβ_42_ monomers and 10% seeds exhibited a sigmoidal increase in bright Aβ_42_ puncta (**Figure 5D**), culminating in a plateau enriched in Aβ_42_ aggregates that stained positive for Amytracker. The seeded treatment at 250 nM Aβ_42_ monomer concentration resulted in slower aggregation, which did not reach a plateau within the observed time even upon seeding (**Figure 5C**). We repeated the seeded aggregation reaction with 500 nM fluorescently-labelled Aβ_42_ monomers in the absence and presence of increasing concentrations of BRICHOS, and monitored the time course aggregation directly on the neurons (**Figure 5E**). The addition of BRICHOS delayed the half-time of the aggregation reaction in a dose-dependent manner (**Figures 5E,F**), in agreement with its effect observed in *in vitro* kinetic assays^28^.

**Figure 5.**
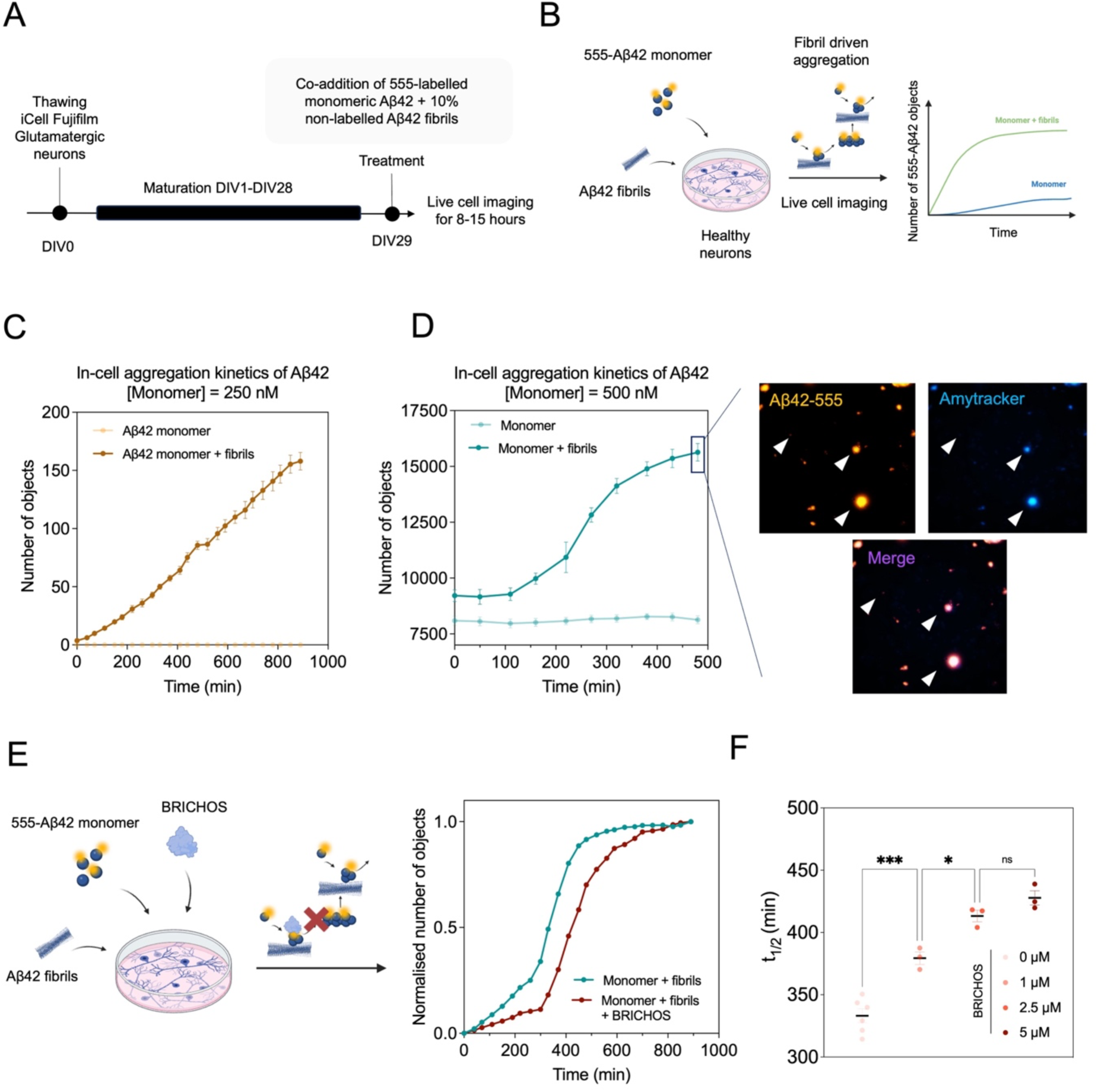
A seeded aggregation kinetics of Aβ_42_ on human glutamatergic neurons can be followed by fluorescently-labelled Aβ_42_ monomers. (**A, B)** Schematic representation of the experimental design and timeline. iCell human glutamatergic neurons were kept in culture for 28 days. On DIV29, neurons were treated with 250 nM or 500 nM of fluorescently labelled Aβ_42_ monomer (Alexa Fluor 555) in the presence or absence of 25 nM or 50 nM respectively of non-labelled Aβ_42_ fibrils. Levels of fluorescent puncta (number of objects) were tracked over time for a period of 8 h to 15 h after the treatment using live cell imaging microscopy. No increase in the number of fluorescent objects were observed for cells treated with monomer alone either at 250 nM **(C)** or 500 nM. **(D).** Neurons treated with seeded monomer at 250 nM showed slow, linear increase in the levels of fluorescent puncta **(C).** Neurons treated with seeded monomer at 500 nM showed a faster, sigmoidal increase in the number of aggregates **(D),** that stained positive for Amytracker in the plateau of the *in-situ* aggregation kinetic reaction. **(E)** Addition of BRICHOS (5 µM) to neurons co-treated with 500 nM of fluorescently-labelled Aβ_42_ and 50 nM of non-labelled fibrils delayed the half-time of the *in situ* aggregation kinetic reaction. **(F)** Effect of increasing the concentration of BRICHOS on the *in-situ* aggregation reaction half-times (t_1/2_) (n = 6 kinetic curves for neurons treated with monomer and fibrils; n = 3 kinetic curves for neurons treated with monomer, fibrils and BRICHOS). Statistical differences were calculated by applying a one-way ANOVA, using the Tukey’s test to correct for multiple comparisons (ns p-value > 0.12, *p-value < 0.033, ***p-value<0.001).

These results show that our method to trigger the *in situ* generation of Aβ42 oligomers can provide quantitative information to assess the effect of potential inhibitors of Aβ_42_ seeded aggregation in the presence of the living cells.

## Discussion

In this study, we developed an approach to study the effects of Aβ_42_ oligomers and aggregates in a cell system. This approach enables the characterisation of the cytotoxic profiles of the Aβ_42_ aggregates formed during the aggregation reaction, and the investigation of the early signs of cell dysfunction caused by Aβ_42_ oligomers. The approach is based on the *in-situ* generation of Aβ_42_ oligomers via secondary nucleation. We obtain this result through the repeated co-addition of Aβ_42_ monomers and low concentrations pre-formed Aβ_42_ fibrils to the cell culture. Our study thus translates into living cells the type of *in vitro* kinetic assays previously used to study the mechanisms governing the aggregation of proteins relevant for neurodegenerative diseases^16,25,26,29^. Unlike the methods that use stabilised Aβ_42_ oligomers^30^, or Aβ_42_ oligomers taken from *in vitro* aggregation reactions^29^, the *in situ* approach avoids the problem of the selection of the aggregate populations to be tested, and overcomes the challenge of isolating metastable Aβ_42_ oligomers^27^. In addition, the exogenous repeated dosage provides control over the level of Aβ_42_ aggregation and severity of disease phenotypes.

The *in situ* protocol enables to study Aβ_42_ aggregate formation in the presence cellular components likely to modulate this process, such as lipid membranes^31–33^, endogenous proteins^34–36^ and metabolites^30,37–39^. Among the differences between the aggregation reaction performed on cell membranes and *in vitro* set-ups, we highlight the greater stochasticity of the former, necessitating fine-tuning of variables such as monomer and seed concentrations, as well as the number of treatments, to develop a robust system for reproducible investigations.

We found that the production levels and the features of the Aβ_42_ oligomers generated in the seeded and unseeded conditions were different. By seeding the aggregation, we produced higher levels of cytotoxic Aβ_42_ oligomers as compared to aggregates generated in the absence of Aβ_42_ fibrils. We also found that Aβ_42_ oligomers and cell-associated aggregates derived from seeded reactions in cells can trigger ROS production in cholinergic-like SH-SY5Y cells, as well as synaptic loss and reduction of the electrophysiological activity in human glutamatergic neurons. These are pathological processes characteristic of early AD stages that make the *in situ* approach suitable for translational studies.

The results reported in this study also illustrate how the level aggregate-derived toxicity is contingent on the cell type used and the chosen readout. We showed how differentiating SH-SY5Y cells to a cholinergic phenotype increases their vulnerability to damage from the aggregates. Similarly, by using iPSC-derived neurons and sensitive readouts, such as electrophysiology-based methods, we could detect dysfunctional effects from aggregate concentrations in the nanomolar range, measured as monomer equivalent, such as those forming in the unseeded aggregation.

We also note that the implementation of this protocol involved several challenges. Aβ_42_ is highly aggregation-prone, making it exceptionally challenging to purify and handle^40^. Additionally, its low content of aromatic amino acids hampers accurate concentration determination, in particular of Aβ_42_ oligomers^27^. Variability in purification from batch to batch can also lead to inconsistent aggregation behaviour, impacting the reproducibility of results across experimental replicates. In practice, all these factors can be responsible of inconsistent results, especially when repeating the assay to compare the efficacy of different candidate compounds in drug discovery pipelines. To overcome these issues, we implemented a series of internal quality controls to ensure the purity of the Aβ_42_ peptide and the maturity of the pre-formed seeds. The reproducibility of the systems and the possibility to measure quantitative parameters, such as the half time of aggregation and amount of aggregates at the plateau phase, allows the characterization of inhibitors of secondary nucleation of Aβ_42_.

We anticipate that the strategy described here can be readily applied to other human iPSC-derived cell types, such as astrocytes or microglia, enabling the implementation of a wide array of functional assays. These studies will enable a broader and more comprehensive exploration of other disease biomarkers linked to Aβ_42_ oligomer production and aggregation, advancing our understanding of AD.

## Conclusions

We reported a protocol to study Aβ_42_ oligomerisation and aggregation via secondary nucleation in cellular systems, and showed that it recapitulated neurite degeneration and synaptic dysfunction in iPSC-derived human glutamatergic neurons. In perspective, we expect that this protocol will serve as a platform for testing compounds that inhibit specific microscopic steps in the Aβ_42_ aggregation process, offering a tool to assess the efficacy of targeted therapies with mechanism of action similar to that of currently approved disease-modifying therapies for AD^16^. We also anticipate that this protocol will be applicable to a wide range of cell types and alternative readouts, as well as to other diseases that involve on-pathway protein aggregation, including Parkinson’s disease, frontotemporal dementia, amyotrophic lateral sclerosis and type 2 diabetes.

## Materials and Methods

### Study design

The objective of this study was to develop an experimental system that recapitulates key hallmarks of AD, particularly: (i) the on-pathway Aβ_42_ aggregation and the generation of the metastable and difficult-to-isolate oligomers in a physiological environment; (ii) the cell dysfunction phenomena associated to these species forming during the on-pathway Aβ_42_ aggregation reaction. To this purpose, we designed a strategy aimed at inducing Aβ_42_ aggregation directly on the living cells by repeating treatments with Aβ_42_ monomers in the presence or absence of pre-formed fibrils. First, we set up experimental conditions in SH-SY5Y cells by optimizing parameters such as Aβ_42_ seed concentrations, Aβ_42_ monomer concentrations, and number of treatments. Secondly, we improved the translational power of the system by correlating the levels of Aβ_42_ aggregation to the synaptic impairment and neurite degeneration in human iPSC-derived glutamatergic neurons. In addition, we confirmed through the use of BRICHOS - a well characterized inhibitor of secondary nucleation - that the aggregation on the neurons occurs via a process of secondary nucleation. Finally, we tested the quantitative power of the cell system by monitoring the aggregation time course in living neurons, obtaining parameters for comparing potential Aβ_42_ aggregation inhibitors.

### Differentiation of SH-SY5Y cells to a cholinergic-like phenotype

Undifferentiated SH-SY5Y (neuroblastoma) cells were subjected to a 7-day protocol adapted from a published protocol^41^. From an 80-90% confluent culture, basal growth medium was replaced by DMEM/F-12, GlutaMAX™ supplemented with 1% v/v of heat-inactivated fetal bovine serum and 10 µM retinoic acid (RA). At days 4 and 7, the growth medium was replaced by DMEM/F-12, GlutaMAX™ supplemented with 1% v/v hiFBS, 10 µM RA and 50 ng/mL of brain-derived neurotrophic factor (BDNF). SH-SY5Y cells were replated into Matrigel-coated flasks or plates after day 8 of differentiation. Serum reduced medium supplemented with RA and BDNF was kept during the duration of all performed experiments.

### Purification and preparation of Aβ_42_ monomers, fibrils and on-pathway aggregates

Recombinant Aβ_42_ (MDAEFRHDSGYEVHHQKLVFFAEDVGSNKGAIIGLMVGGVVIA) was expressed and purified, and monomers were prepared as previously reported^31,40^. In brief, *E. coli* cells were sonicated, followed by the solubilization of inclusion bodies using 8 M urea. This was succeeded by ion exchange chromatography using diethylaminoethyl cellulose resin in a batch mode and subsequent lyophilization. Samples were further purified using a Superdex 75 HR 26/60 column (GE Healthcare, Buckinghamshire, UK). The eluted fractions were then subjected to SDS-PAGE to check for the target protein. Fractions containing the recombinant protein were pooled, quickly frozen in liquid nitrogen, and then lyophilized once more.

Solutions of monomeric peptides either to generate on-pathway aggregates, fibrils or to be used directly for cell treatments were prepared as follows. Lyophilised Aβ_42_ was dissolved in 6 M guanidinium hydrocholoride (GuHCl). Monomeric forms were purified from potential oligomeric species and salt using a Superdex 75 10⁄300 GL column (GE Healthcare) at a flowrate of 0.7 mL/min, and were eluted in 20 mM sodium phosphate buffer, pH 8. Importantly, no EDTA or NaN_3_ was supplemented in the buffer, to prevent cell toxicity. Protein elution was tracked with the use a chromatogram, assessing both the elution volume and absorbance levels at λ_280_. The centre of the peak was collected, using pre-chilled protein LoBind Tubes. The concentration of Aβ_42_ was determined from the absorbance of the integrated peak area (ε = 1490 M^-1^ cm^-1^ and MW = 4461 g/mol).

To prepare kinetic experiments, the monomer was diluted up to the desired concentration using the same buffer (20 mM sodium phosphate buffer, pH 8, without EDTA and NaN_3_). 20 µM of thioflavin T (ThT) was added from a 1 mM stock to some samples to follow the aggregation kinetic reaction. 80 µL of each monomer sample was pipetted into at least three wells of a 96-well half-area, low-binding, clear bottom and PEG-coated plate (Corning 3881). T_0_, t_lag_ and t_half_ samples (without ThT) were isolated and transferred to fresh protein LoBind Tubes prior cell treatment.

For the generation of Aβ_42_ fibrils, freshly purified monomeric protein was filtered using a 0.22 μm low protein binding filter unit (#SLGV004SL, Millex ®-GV) in a final volume 500 μL of 20 μM concentration. Filtered monomeric peptides were kept in 2 mL LoBind Tubes (#0030108450, Eppendorf) at 37 °C for 72 h, without shaking. ThT-binding assay was performed for every batch to assess reproducibility in the average cross-β content of the fibrillar species.

### Purification of BRICHOS

The recombinant BRICHOS domain pro-SP-C was purified as previously reported^42^. In brief, the BRICHOS construct was transformed into *E. coli* Origami (DE3) pLysS. Bacteria were grown in LB medium with 100 μg/ml ampicillin at 30°C for 16h. Protein expression was induced with 0.25 mM IPTG, followed by incubation at 25°C for 6h. Cells were harvested, resuspended in 20 mM NaP buffer containing 5 mM imidazole, pH 7.0, and stored at -80°C. For protein purification, bacterial pellets were sonicated and centrifuged. The filtered supernatant was applied to a Ni-NTA column pre-equilibrated with 20 mM NaP, 5 mM imidazole, pH 7.0. The column was washed with an imidazole gradient (5-50 mM) and the protein was eluted with 100 mM imidazole in 20 mM NaP, pH 7.0. After dialysis in 20 mM NaP, pH 7.0 to remove imidazole, the His6 and thioredoxin tags were removed using thrombin treatment from bovine plasma, followed by another Ni-NTA column passage to separate the tags.

### Expression, purification and labelling of the S8C-Aβ_42_ variant with Alexa Fluor 555

A panel of single cysteine variants for the peptide Aβ_42_ was previously reported^43^. Given the similarities in its aggregation kinetics behaviour and the morphology of the derived fibrils with the wild-type isoform, the single cysteine variant S8C was employed in the present study. In brief, the plasmid carrying the synthetic gene Aβ_42_ S8C was obtained from Prof. Sara Linse (Lund University) and transformed in the of *E. coli* strain BL21 DE3 pLysS star. The peptide was expressed in auto-induction medium. Aβ_42_ was then purified using ion exchange chromatography, using 50 mM NaCl for elution, followed by size exclusion chromatography (SEC) on a 26 × 600 mm Superdex 75 column. Buffers for the purification were supplemented with 1 mM dithiothreitol (DTT) to prevent the cysteine mutants to dimerise. DTT was, however, removed for the final SEC. Purified monomers were lyophilised and further resuspended in Milli-Q water to reach a final concentration of approximately 14 µM in 50 µL of volume. A final concentration of 3 to 4 mM of Alexa Fluor 555 was added to the dissolved peptide. The mixture was incubated overnight at 4 °C. The mixture was then added to 1 mL of 6 M GuHCl prepared in 20 mM sodium phosphate pH 8.5. Labelled monomer was purified via a SEC using a Superdex 75 10/300 column in 20 mM sodium phosphate pH 8. To follow the purification of the labelled peptide, together with the remaining excess of dye and unlabelled protein, the absorbance at both 280 nm and 555 nm was monitored using a Quadtech detector. The concentration of the labelled peptides was determined using the correction factor for Alexa Fluor 555, following an equation previously reported.

### iCell GlutaNeurons culture and maintenance

iPSC-derived iCell GlutaNeurons (iGNCs) were obtained from Cellular Dynamics International and cultured according to manufacturer instructions.

### Preparation of cells for immunocytochemistry assays

For immunocytochemistry assays, either Corning^®^ 96 Well TC-Treated Microplates or PerkinElmer 96 Well Cell Carrier Ultra plates were coated with 80 µL per well of a 0.07% PEI solution. Plates were incubated at 37 °C for 1 h before rinsing the content twice with 260 µL/well of sterile D-PBS (+/+) (#14040141, Corning). Plates were left to dry overnight at room temperature. Wells were then coated with a 1X solution of Matrigel diluted in cold DMEM. Plates were incubated at 37 °C for 1 h prior plating. iGNCs were thawed as per manufacturer’s instructions and plated at a density of 80k cells/well in complete BrainPhys medium (95 mL of BrainPhys Neuronal Medium supplemented with 2 mL of iCell Neural Supplement, 1 mL of iCell Nervous System Supplement, 1 mL of N-2 Supplement, 1 µg/mL of laminin 2020 (L2020) and 1 mL of penicillin-streptomycin. Medium was replaced following recommended regimens until cell treatment at DIV31 and DIV34.

### Preparation of cells for multi-electrode arrays (MEA)

For electrophysiology assays, 96-well MEA plates (Axion Biosystems) were coated with PEI as per manufacturer’s indication. iGNCs were thawed and concentrated at a density of 12 M cells/mL in a dotting solution composed by complete BrainPhys medium supplemented with a 10% v/v of L2020. 11 µL were spotted at the centre of each well. Cells were incubated at 37 °C for 1 h prior to the addition of 300 µL/well of complete BrainPhys medium.

### Multiple treatment on human SH-SY5Y cells

Freshly differentiated human SH-SY5Y cells were plated in Corning^®^ 96 Well TC-Treated Microplates coated with Matrigel at a density of 30k cells/well in 100 µL/well of DMEM/F-12, GlutaMAX™ supplemented with 1% hiFBS, 10 µM RA and 50 ng/mL of BDNF. Cells were incubated at 37 °C for 24 h prior treatment. On the day of the first treatment, 50 µL of medium per well was removed and replaced by 50 µL of each treatment solution, composed by 25 µL of growth medium and 25 µL of a mixture of 20 µL of 2 µM Aβ_42_ and 5 µL of 200 nM of pre-formed amyloid fibrils both diluted in 20 mM NaP buffer pH 8. Every 24 h or 48 h, cells were either fixed or treated again. For each treatment, 50 µL of medium per well was collected for measurement of oligomer secretion and replaced by 50 µL of each treatment solution.

### Multiple treatment on iCell GlutaNeurons

For cells cultured in 96-well plate formats: At DIV29, full medium was removed and 100 µL of fresh complete BrainPhys medium was added per well. 100 µL of treatment solution was prepared by mixing 75 µL of BrainPhys medium, 20 µL of 5µM monomeric Aβ_42_ and 5 µL of 500 nM pre-formed fibrils when appropriate. Fibrils were always added after homogenising the monomer with the medium. Solution was carefully mixed 8 times with the use of a pipette before treatment. For BRICHOS treatment, 48 µL of BrainPhys medium, 20 µL of 5 µM of Aβ_42_ monomer and 1 µL of 10 µM BRICHOS were carefully mixed 20 times with the use of a pipette. The mixture was incubated at room temperature for 30 min. Afterwards, 5 µL of 500 nM pre-formed fibrils were added, and the solution was homogenised by pipetting 8 times before treatment. 48 h after the first treatment, 100 µL of medium from treated cells was transferred to a mirror 96-well plate, flash-frozen and stored at -80 °C prior oligomer content analysis. The second treatment was prepared as explained above. After 48 h, supernatant was collected, and cells were fixed for immunostaining.

### For cells cultured in 96-well MEA plates

Six hours before the treatment, full medium was transferred to an empty, sterile 96-well plate. 100 μL of conditioned medium was transferred back to the MEA plate wells and 100 μL of fresh complete BrainPhys medium was added per well. After 6h, 100 μL per well were collected for treatment preparation. For the treatment, 80 μL of conditioned medium were mixed with 18 μL of 5 μM of monomeric Aβ42 and 2 μL of 0.5 μM pre-formed fibrils when appropriate. The mixture was carefully mixed 8 times with the use of a pipette before cells treatment. 9-10 wells were treated per condition, using 100 μl/well of the medium, Aβ and seeds mixtures. The same procedure was repeated again for the subsequent treatments.

### Capture-detector ELISA for quantification of secreted Aβ_42_ oligomers

Streptavidin-coated 96-well plates were washed with 300 µL per well of 0.05% Tween-20 in PBS. Afterwards, each well was coated with 70 µL of a 1:500 dilution of a biotinylated 6E10 antibody (#803008, BioLegend) in 0.05% Tween-20. Capture antibody was incubated for 1 h at RT under shaking conditions (250 rpm). Cell supernatants were thawed at RT and centrifuged at 4000 rpm for 15 min at 4 °C to remove insoluble aggregates and cellular debris. Protease inhibitor solution was added at 1X final concentration to each sample (COEDTAF-RO, Roche). Capture antibody was washed twice with 0.05% Tween-20 in PBS before sample addition. Samples were incubated for 2 to 3 h at 4 °C under shaking conditions (250 rpm). The plates were then washed three times with TBS prior addition of HRP-6E10 detector antibody (#803012, BioLegend) diluted in a 5% BSA solution in TBS at a concentration of 0.1 µg/mL. Detector antibody was incubated 1 h at RT under agitation (250 rpm). Plates were washed three times in TBS. Afterwards, 100 µL of TMB ELISA substrate was added per well. The reaction was left to develop for 30 min in the dark after being stopped with the addition of 50 µL per well of 2M HCl. Absorbance values were measured at a wavelength of 450 nm using a plate reader (BMG Labtech, Aylesbury, UK).

### Immunocytochemistry and staining for Aβ_42_ amyloid aggregates

Cells were fixed with a solution of 4% PFA diluted in D-PBS (+/+), at room temperature for 10 min. Fixed cells were then washed three times with D-PBS (+/+), followed by permeabilization with 0.1% v/v of Triton X-100 in D-PBS (+/+) (#85111, Thermo Scientific) for 30 min at RT. Afterwards, cells were treated with a blocking solution containing 2% BSA, 3% goat serum, in D-PBS (+/+) for 1 h at RT. Blocking solution was removed and cells were incubated with primary antibodies in blocking solution overnight at 4 °C. After four washes with D-PBS (+/+), cells were then incubated with Alexa Fluor-conjugated secondary antibodies for 1 h at room temperature in the dark. Cells were wash other four times with D-PBS (+/+). Fibrillar aggregates were stained using the amyloid-binding dye Amytracker 638 (Ebba Biotech) at 1:500 dilution in D-PBS (+/+). Nuclei was stained using Hoechst 33342 (#H3570, Thermo Fisher) at 1:1000 dilution, also in D-PBS (+/+).

### Immunostaining antibodies

The following primary and secondary antibodies were used for immunostaining experiments: Mouse anti-Aβ/APP antibody (1:500, SigmaAldrich, #MABN10); mouse anti-synapsin 1 (1:1000, Synaptic Systems, #106011), rabbit anti-β-III-catenin (1:1000, Abcam, #ab52623); Alexa Fluor 555 goat anti-Mouse IgG (H+L) (1:1000, Thermo Fisher, #A21424), Alexa Fluor 488 goat anti-Rabbit IgG (H+L) (1:1000, Thermo Fisher, # A11001).

### Electrophysiology

The spontaneous electrical network activity was recorded 24 h before the first treatment (DIV40), 48 h after the first treatment (DIV42) and 72 h after the second treatment (DIV47) using the Axion Maestro Pro system. During the recording procedure, a 20-min equilibration period was allowed, followed by one measurement of 10 min. All recordings were conducted using the Axion Integrated Study (AxIS) software, under a controlled environment of 37 °C and 5% CO_2_. The electrical activity was measured with a gain of 1000x and a sampling frequency of 12.5 kHz. Before spike detection, a Butterworth band-pass filter ranging from 100 to 3000 Hz was applied. Spike detection was performed using the AxIS adaptive spike detector, with a threshold set at 6 times the root mean square (RMS) noise on each electrode. An electrode was considered active if it exhibited a spike rate of at least 5 spikes per min. These recording and analysis parameters were employed to evaluate and characterize the spontaneous electrical network activity in the experimental setup. Each recording was re-recorded using the AxIS software, generating .spk files. Network bursts were identified using the Axion Neural Metric Tool. These detection methods yielded 57 parameters related with individual neuronal activity, network burst and network synchrony. Three representative parameters were chosen as representative of the effect of the aggregates derived from exogenous feeding with monomer and fibrils on electrophysiological activity.

## Author contributions

A.G.D., E.S., B.M., S.C. and M.V. designed research; A.G.D., and E.S., performed research; S.C., J.M., G.S., R.C. and K.Y., prepared and purified protein; K.Y., prepared and labelled protein. Y.B., contributed with microscopy results. A.G.D., E.S., A.P., I.K-S. and B.M., analysed data. A.G.D., G.A.U, B.M., and M.V., wrote the paper.

## References

1 Jack Jr, C. R., et al. Revised criteria for diagnosis and staging of Alzheimer’s disease: Alzheimer’s association workgroup. Alzheimers Dement. (2024).

2 Salvadó, G. et al. Disease staging of Alzheimer’s disease using a CSF-based biomarker model. *Nat*. Aging 4, 694–708 (2024).

3 Therriault, J. et al. Biomarker-based staging of Alzheimer disease: Rationale and clinical applications. Nat. Rev. Neurol. 20, 232–244 (2024).

4 Selkoe, D. J. & Hardy, J. The amyloid hypothesis of Alzheimer’s disease at 25 years. EMBO Mol. Med. 8, 595–608 (2016).

5 Knopman, D. S. et al. Alzheimer disease. Nat. Rev. Dis. Primers 7, 33 (2021).

6 Hampel, H. et al. The amyloid-β pathway in Alzheimer’s disease. Mol. Psychiatry 26, 5481–5503 (2021).

7 Lambert, M. P. et al. Diffusible, nonfibrillar ligands derived from Aβ1–42 are potent central nervous system neurotoxins. Proc. Natl. Acad. Sci. USA 95, 6448–6453 (1998).

8 Haass, C. & Selkoe, D. J. Soluble protein oligomers in neurodegeneration: Lessons from the Alzheimer’s amyloid β-peptide. Nat. Rev. Mol. Cell Biol. 8, 101–112 (2007).

9 Knowles, T. P., Vendruscolo, M. & Dobson, C. M. The amyloid state and its association with protein misfolding diseases. Nat. Rev. Mol. Cell Biol. 15, 384–396 (2014).

10 Rinauro, D. J., Chiti, F., Vendruscolo, M. & Limbocker, R. Misfolded protein oligomers: Mechanisms of formation, cytotoxic effects, and pharmacological approaches against protein misfolding diseases. Mol. Neurodegener. 19, 20 (2024).

11 De, S. et al. Different soluble aggregates of Aβ42 can give rise to cellular toxicity through different mechanisms. Nat. Comm. 10, 1541 (2019).

12 Mannini, B. et al. Toxicity of protein oligomers is rationalized by a function combining size and surface hydrophobicity. ACS Chem. Biol. 9, 2309–2317 (2014).

13 Selkoe, D. J. The advent of Alzheimer treatments will change the trajectory of human aging. *Nat*. Aging 4, 453–463 (2024).

14 Cummings, J. et al. Alzheimer’s disease drug development pipeline: 2023. Alzheimer’s & Dementia 9, e12385 (2023).

15 Sevigny, J. et al. The antibody aducanumab reduces Aβ plaques in Alzheimer’s disease. Nature 537, 50–56 (2016).

16 Linse, S. et al. Kinetic fingerprints differentiate the mechanisms of action of anti-Aβ antibodies. Nat. Struct. Mol. Biol. 27, 1125–1133 (2020).

17 Van Dyck, C. H. et al. Lecanemab in early Alzheimer’s disease. N. Engl. J. Med. 388, 9–21 (2023).

18 Cummings, J. et al. Lecanemab: Appropriate use recommendations. J. Prev. Alzheimers Dis. 10, 362–377 (2023).

19 Mintun, M. A. et al. Donanemab in early Alzheimer’s disease. N. Engl. J. Med. 384, 1691–1704 (2021).

20 Cummings, J., Osse, A. M. L., Cammann, D., Powell, J. & Chen, J. Anti-amyloid monoclonal antibodies for the treatment of Alzheimer’s disease. BioDrugs 38, 5–22 (2024).

21 Cohen, S. I. et al. Proliferation of amyloid-β42 aggregates occurs through a secondary nucleation mechanism. Proc. Natl. Acad. Sci. USA 110, 9758–9763 (2013).

22 Michaels, T. C. et al. Dynamics of oligomer populations formed during the aggregation of Alzheimer’s Aβ42 peptide. Nat. Chem. 12, 445–451 (2020).

23 Cohen, S. I., Vendruscolo, M., Dobson, C. M. & Knowles, T. P. From macroscopic measurements to microscopic mechanisms of protein aggregation. J. Mol. Biol. 421, 160–171 (2012).

24 Meisl, G. et al. Molecular mechanisms of protein aggregation from global fitting of kinetic models. Nat. Protoc. 11, 252–272 (2016).

25 Habchi, J. et al. Systematic development of small molecules to inhibit specific microscopic steps of Aβ42 aggregation in Alzheimer’s disease. Proc. Natl. Acad. Sci. USA 114, E200–E208 (2017).

26 Chia, S. et al. SAR by kinetics for drug discovery in protein misfolding diseases. Proc. Natl. Acad. Sci. USA 115, 10245–10250 (2018).

27 Kulenkampff, K., Wolf Perez, A.-M., Sormanni, P., Habchi, J. & Vendruscolo, M. Quantifying misfolded protein oligomers as drug targets and biomarkers in Alzheimer and parkinson diseases. Nat. Rev. Chem. 5, 277–294 (2021).

28 Cohen, S. I. et al. A molecular chaperone breaks the catalytic cycle that generates toxic Aβ oligomers. Nat. Struct. Mol. Biol. 22, 207–213 (2015).

29 Habchi, J. et al. An anticancer drug suppresses the primary nucleation reaction that initiates the production of the toxic Aβ42 aggregates linked with Alzheimer’s disease. Sci. Adv. 2, e1501244 (2016).

30 González Díaz, A., Cataldi, R., Mannini, B. & Vendruscolo, M. Preparation and characterization of zn (ii)-stabilized Aβ42 oligomers. ACS Chem. Neurosci. (2024).

31 Habchi, J. et al. Cholesterol catalyses Aβ42 aggregation through a heterogeneous nucleation pathway in the presence of lipid membranes. Nat. Chem. 10, 673–683 (2018).

32 Sanguanini, M. et al. Complexity in lipid membrane composition induces resilience to Aβ42 aggregation. ACS Chem. Neurosci. 11, 1347–1352 (2020).

33 Baumann, K. N. et al. A kinetic map of the influence of biomimetic lipid model membranes on Aβ42 aggregation. ACS Chem. Neurosci. 14, 323–329 (2022).

34 Arosio, P. et al. Kinetic analysis reveals the diversity of microscopic mechanisms through which molecular chaperones suppress amyloid formation. Nat. Comm. 7, 10948 (2016).

35 Xia, Z. et al. Co-aggregation with apolipoprotein e modulates the function of amyloid-β in Alzheimer’s disease. Nat. Comm. 15, 4695 (2024).

36 Cristóvão, J. S. et al. The neuronal s100b protein is a calcium-tuned suppressor of amyloid-β aggregation. Sci. Adv. 4, eaaq1702 (2018).

37 Abelein, A., Gräslund, A. & Danielsson, J. Zinc as chaperone-mimicking agent for retardation of amyloid β peptide fibril formation. Proc. Natl. Acad. Sci. USA 112, 5407–5412 (2015).

38 Mannini, B. et al. Stabilization and characterization of cytotoxic Aβ40 oligomers isolated from an aggregation reaction in the presence of zinc ions. ACS Chem. Neurosci. 9, 2959–2971 (2018).

39 Chia, S. et al. A relationship between the structures and neurotoxic effects of Aβ oligomers stabilized by different metal ions. ACS Chem. Neurosci. (2024).

40 Walsh, D. M. et al. A facile method for expression and purification of the Alzheimer’s disease-associated amyloid β-peptide. FEBS J. 276, 1266–1281 (2009).

41 de Medeiros, L. M. et al. Cholinergic differentiation of human neuroblastoma SH-SY5Y cell line and its potential use as an in vitro model for Alzheimer’s disease studies. Mol. Neurobiol. 56, 7355–7367 (2019).

42 Willander, H. et al. BRICHOS domains efficiently delay fibrillation of amyloid β-peptide. J. Biol. Chem. 287, 31608–31617 (2012).

43 Thacker, D., Bless, M., Barghouth, M., Zhang, E. & Linse, S. A palette of fluorescent Aβ42 peptides labelled at a range of surface-exposed sites. Int. J. Mol. Sci. 23, 1655 (2022).

44 Michno, W. et al. Following spatial Aβ aggregation dynamics in evolving Alzheimer’s disease pathology by imaging stable isotope labeling kinetics. Sci. Adv. 7, eabg4855 (2021).

