## Supplementary Information for "*In situ* generation of Aβ_42_ oligomers via secondary nucleation triggers neurite degeneration and synaptic dysfunction in human iPSC-derived glutamatergic neurons"

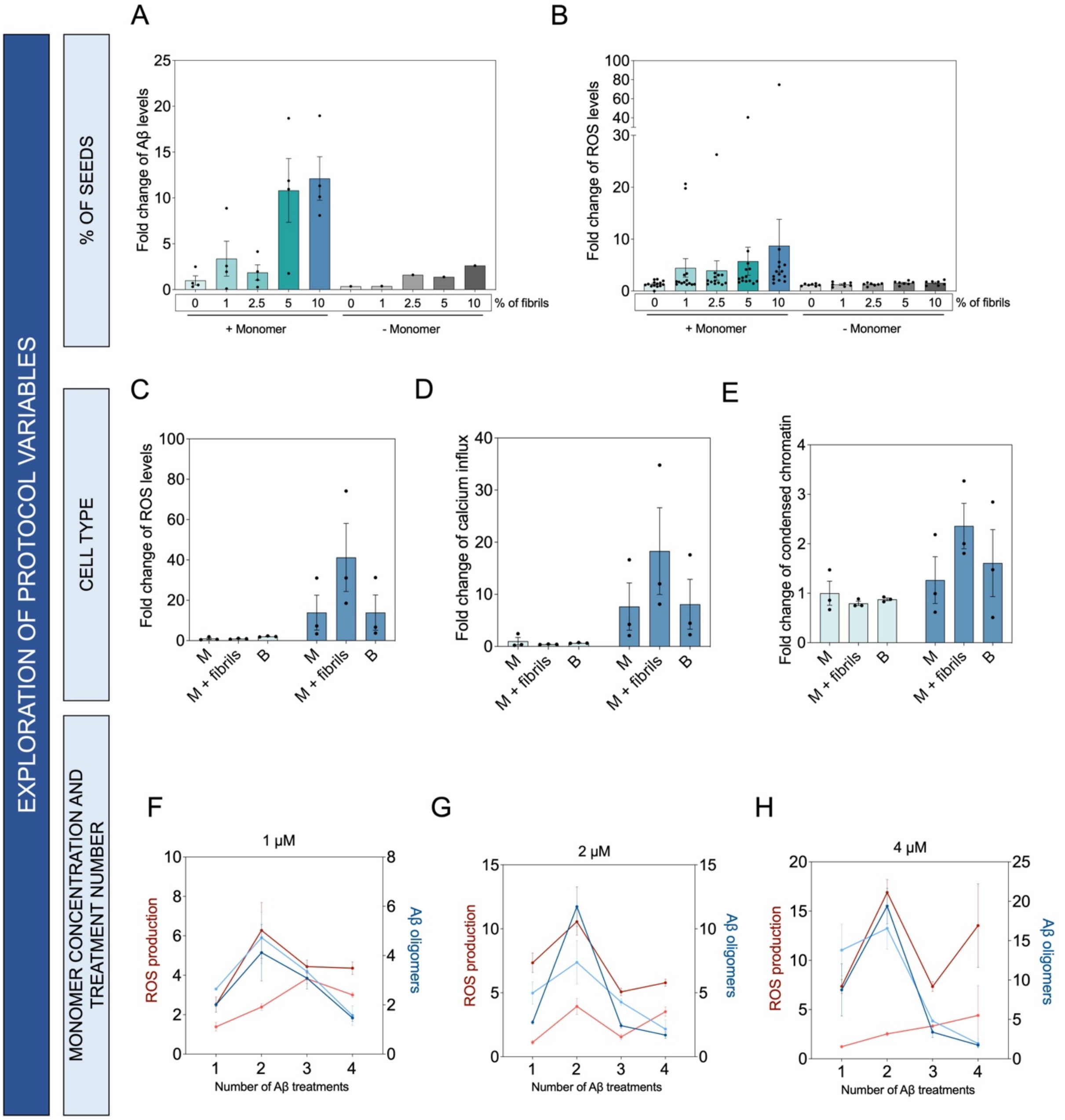


**Figure S1. Optimization of the *in situ* protocol to generate on-pathway Aβ42 oligomers.** We optimized the three main components of the protocol: percentage of seeds (A,B), cell type and monomer concentration (C-E), and treatment number (F-H). **(A,B)** Non-differentiated human SH-SY5Y cells were co-treated with 500 nM of freshly-purified Aβ42 monomers and 5 nM (1%), 12.5 nM (2.5%), 25 nM (5%) and 50 nM (10%) of pre-formed fibrils (monomer equivalents). Cells were either immunoassayed using W02 to quantify the total Aβ levels (A) and stained with CellRox to assess ROS production (B). Aβ42 levels were calculated by quantifying the total area of fluorescent signal derived from W02. ROS production was calculated by quantifying the total fluorescence derived from the CellRox dye. Data were normalised by the total amount of cells (total bright field area). The fold change was calculated over the fluorescence derived from the cells treated with only with monomer (M + 0%) (n = 4 for Aβ42 immunostaining and n = 7 or 12 for ROS quantification). **(C-E)** Non-differentiated and cholinergic-like SH-SY5Y cells were treated with 500 nM of A$\beta_{42}$ monomer (M), 500 nM of A$\beta_{42}$ monomer and 5 nM of pre-formed fibrils (M + seeds) or buffer. 24 h after the treatment, cells were stained with CellRox to measure ROS production (C), Fluo4 to measure intracellular calcium levels (D), and Hoechst 33342 to measure chromatin condensation (E). The total fluorescence derived from each dye was normalised by total amount of cells (total bright field area). The fold change was calculated over the fluorescence derived from non-differentiated monomer treated cells (n = 3). **(F-H)** Cholinergic-like SH-SY5Y cells were treated every 24 h for 4 consecutive days with 1 μM A$\beta_{42}$ monomers and 100 nM of pre-formed A$\beta_{42}$ fibrils (seeds) (F), 2 μM monomeric A$\beta_{42}$ and 200 nM of pre-formed A$\beta_{42}$ fibrils (seeds) (G), or 4 μM monomeric A$\beta_{42}$ and 400 nM of pre-formed A$\beta_{42}$ fibrils (seeds) (H). 24 h after each treatment, soluble A$\beta_{42}$ aggregates were quantified from the supernatant of treated cells by a 6E10-6E10 homotypic ELISA and, concurrently, cells were stained with CellRox to assess ROS production via live-cell imaging. Data are reported as the fold change of the total absorbance (A$\beta_{42}$ oligomer ELISA) and normalised fluorescence (ROS) over the cells treated only once with buffer (n = 5).

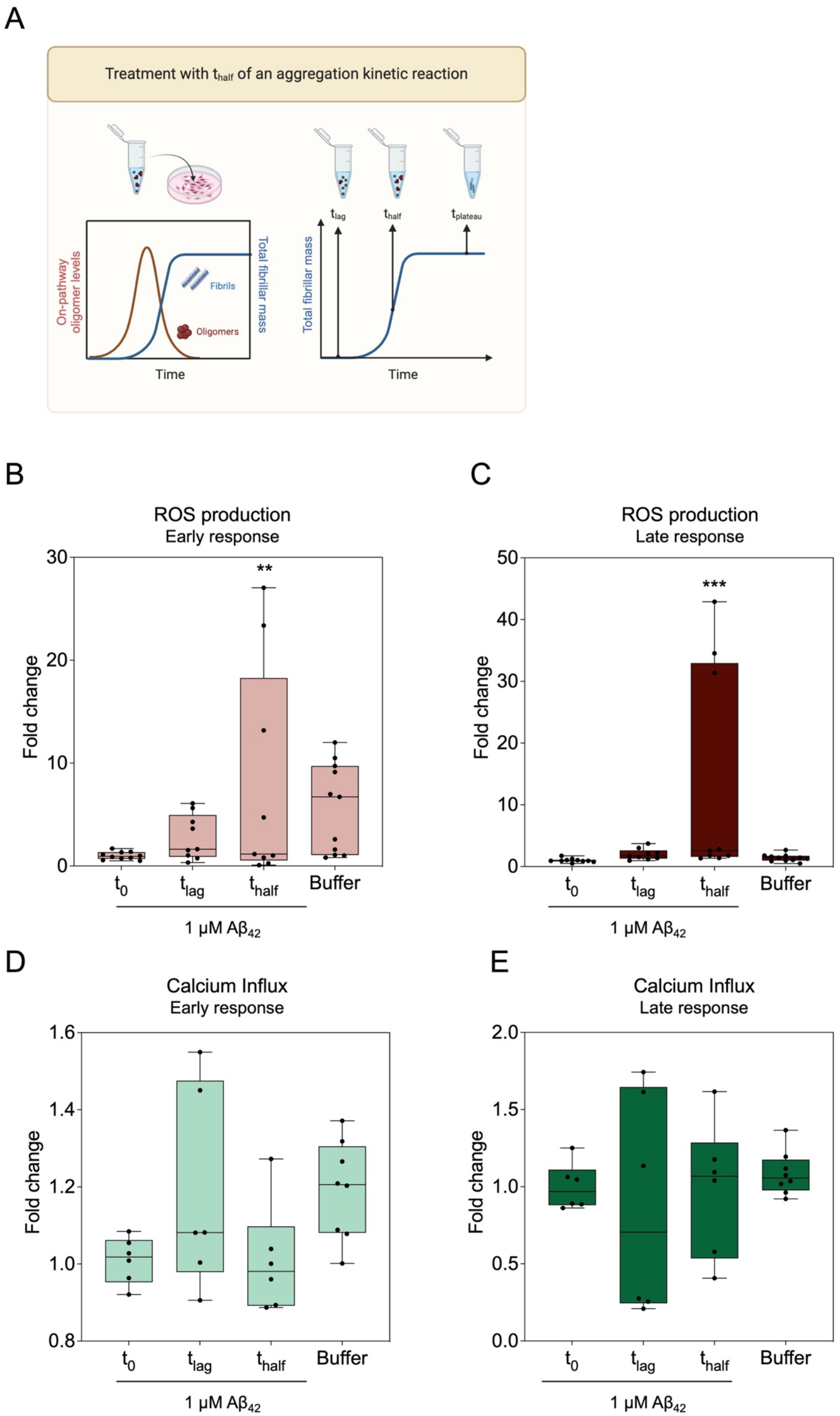
A
Treatment with $t_{half}$ of an aggregation kinetic reaction
On-pathway oligomer levels
Total fibrillar mass
Fibrils
Oligomers
Time
$t_{lag}$
$t_{half}$
$t_{plateau}$
Total fibrillar mass
Time
B
ROS production
Early response
Fold change
**
$t_0$ $t_{lag}$ $t_{half}$ Buffer
1 μM $Aβ_{42}$
C
ROS production
Late response
Fold change
***
$t_0$ $t_{lag}$ $t_{half}$ Buffer
1 μM $Aβ_{42}$
D
Calcium Influx
Early response
Fold change
$t_0$ $t_{lag}$ $t_{half}$ Buffer
1 μM $Aβ_{42}$
E
Calcium Influx
Late response
Fold change
$t_0$ $t_{lag}$ $t_{half}$ Buffer
1 μM $Aβ_{42}$

**Figure S2. On-pathway Aβ$_{42}$ aggregates derived from *in vitro* Aβ$_{42}$ aggregation kinetics show variable cytotoxic effects in SH-SY5Y cells.** (**A**) SH-SY5Y cells were treated with 1 µM (monomer equivalents) of pre-aggregated Aβ$_{42}$ for 0 min ($t_0$), 10 min ($t_{lag}$) or 40 min ($t_{half}$). We assessed the cytotoxic effect solely derived from the buffer by treating cells with the same percentage of 20 mM sodium phosphate without any protein. (**B-E**) Levels of ROS (**B, C**) or calcium influx (**D, E**) were measured immediately (early response) or 24 h after the treatment (late response) via live-cell imaging using the CellRox™ or Fluo4™ dyes. For both readouts, the total fluorescence derived from CellRox™ and Fluo4™ was calculated and normalised by the total amount of cells (total bright field area). The fold change of the normalised fluorescence was calculated over the cells treated with $t_0$. Statistical differences were calculated by applying one-way ANOVA analysis, using the Dunnett's test to correct for multiple comparisons (N = 3 for CellRox (**p-value < 0.01, ***p -value < 0.001 and N = 2 for calcium influx).

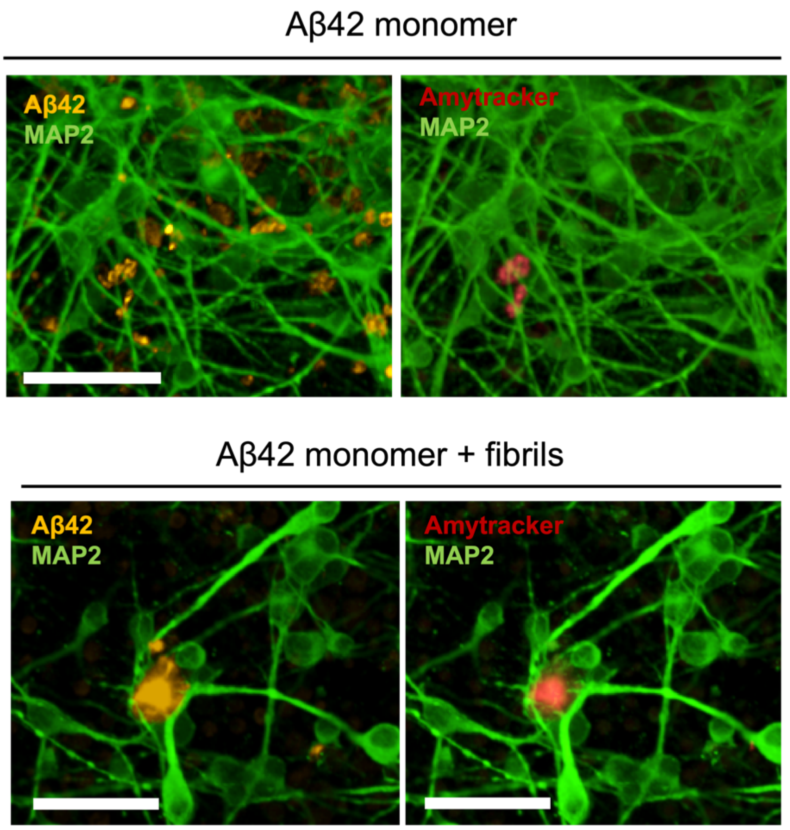


**Figure S3. Morphology of Aβ$_{42}$ aggregates differ between non-seeded or seeded conditions.** Representative pictures of Aβ$_{42}$ aggregates from glutamatergic neurons treated twice with 500 nM of Aβ$_{42}$ monomers and 50 nM of preformed Aβ$_{42}$ fibrils. The antibody W02 was used to detect Aβ$_{42}$ by immunocytochemistry. Amytracker was used to stain for fibrillar aggregates. Scale bar = 50 µm.
